# Direct laser writing of 3D electrodes on flexible substrates

**DOI:** 10.1101/2022.06.07.495165

**Authors:** Morgan A. Brown, Kara M. Zappitelli, Loveprit Singh, Rachel C. Yuan, Melissa A. Bemrose, Valerie Brogden, David J. Miller, Stuart F. Cogan, Timothy J. Gardner

## Abstract

This report describes a 3D microelectrode array integrated on a thin-film flexible cable for neural recording in small animals. The micro electrode array fabrication process integrates traditional silicon thin-film processing techniques and direct laser writing of 3D structures at micron resolution via two-photon lithography. While direct laser writing of 3D printed electrodes has been described before, this report is the first to provide a method for high-aspect-ratio laser-written structures integrated with microfabricated electrical traces. One prototype is a 16-channel array composed of 350 μm long shanks spaced on a grid with 90 μm pitch. Other devices shown here include biomimetic mosquito-needles that penetrate through the dura of birds and porous electrodes designed to promote tissue ingrowth or enhance charge injection capacity for neural stimulation. These devices are just a few examples of a new design space that will enable high-channel-count 3D electrode arrays with features definable at single micrometer resolution. Using a custom laser writer, the 3D printing process is rapid (1 mm^3^/min). This high-speed printing combined with standard wafer-scale processes will enable efficient device fabrication and new studies examining the relationship between electrode geometry and electrode performance. We anticipate highest impact in small animal models, nerve interfaces, retinal implants, and other applications requiring small, high density 3D electrodes.

## Introduction

New technologies are required for high-density chronic neural recording in animal studies and human clinical devices. While a number of promising electrode technologies have emerged in recent years, the devices with the highest impact are those that can be manufactured efficiently. Typically, this means devices built in a cleanroom via thin-film processes developed for the integrated circuit industry. Examples of electrodes in this class include polymer electrodes developed at Lawrence Livermore Labs [1] and Neuralink Inc. [2], silicon electrodes developed in Michigan and Neuronexus Inc. [3], and the transformative, high-density silicon electrodes designed by Neuropixel [4]. While cleanroom fabrication methods provide the required miniaturization and scalability, thin-film devices are by nature, planar. Three dimensional (3D) electrode structures, such as the “bed of nails” design, have historically been assembled by hand, though microfabricated electrodes have been folded into 3D shapes for in-vitro applications [5]. The most widely used 3D electrode array, the Utah array, involves mechanical cutting or dicing of silicon. This step requires rigid backing and the minimum spacing is limited by the width of the cutting tool [6]. As a result, Utah arrays are too large to be used in small animals such as mice and songbirds, and their relatively large shanks also limit chronic performance due to foreign body tissue responses. The Plexon N-Form (3D) array has similar physical size constraints, prohibiting many applications in small animals, nerves or retina.

3D-printed electrodes provide a new alternative to current electrode designs. Recent devices developed at Carnegie Mellon University have demonstrated the concept of 3D-printed bed-of-nails electrodes using an aerosol jet conformal printing method [7]. Though groundbreaking, the devices described in a pre-print are limited by low resolution (10 μm) in the aerosol jet process.

In this work, we describe a high-resolution 3D electrode array fabricated with a combination of two-photon lithography and thin-film fabrication processes that can both achieve micron resolution. While multielectrode arrays fabricated on silicon via two-photon lithography were reported over a decade ago, the fabrication steps utilized either nanoimprint or proximity-mask photolithography, limiting 3D structures to a maximum height of a few micrometers. Furthermore, the previous devices were exclusively fabricated on rigid glass or Si substrates [8–10]. Here we report a fabrication process yielding high aspect-ratio devices (>10:1) integrated on flexible polyimide or parlyene C films, in a form factor suitable for chronically implanted devices. The 3D printing process described here is flexible, allowing the creation of distinct height profiles along the electrode array and various electrode shapes – fully customizable electrode arrays that conform to specific anatomical features of the brain.

## Results

### High-speed custom 3D printer enables efficient fabrication

Two-photon lithography is a 3D printing method that uses femtosecond pulses of infrared light to polymerize an ultraviolet photoresist at the focal point of a high-numerical-aperture lens. By changing the position of the focal point within the liquid photoresist, complex polymer shapes can be written at micron resolution. We recently described an open-source 3D printing system that uses a resonant scan mirror to increase printing speeds by 1-2 orders of magnitude [11] relative to commercially-available galvanometer-based printers. In this system, a design that fills a cubic millimeter of space at near micron resolution takes approximately one minute to print. The printer incorporates fluorescence imaging and reflected-light-sensing pathways that provide real-time information about the degree of polymer crosslinking as well as surface localization. Surface localization to within a micron during the print process is a critical requirement to achieve strong adhesion between the 3D print and the substrate.

### 3D printed arrays for neural recording

The prototype device is a 16-channel array composed of 350 μm tall electrodes with a 20 μm diameter at the recording tip. The electrodes are spaced at 90 μm, and the device includes a hybrid polyimide/parylene flex cable running from the electrodes to an external connector.

The fabrication schematic in **figure 2** outline a simplified geometry, which does not include the 16 mm long polyimide ribbon cable shown in **figure 1**. The first five steps in **figure 2** are standard steps in wafer-scale thin-film processing. To perform the 3D prints in step 6, the wafer is transferred to the 3D printer and liquid photopolymer is applied. The microscope objective is immersed in this liquid photopolymer, and the 3D shapes are printed. Each 16-channel array of polymer spikes is printed in approximately 10 minutes using the 3D printer’s integrated surface finding and substrate positioning features. Following the print process, the wafer and 3D components are metallized simultaneously via a non-directional sputter deposition process.

**Figure 1.**
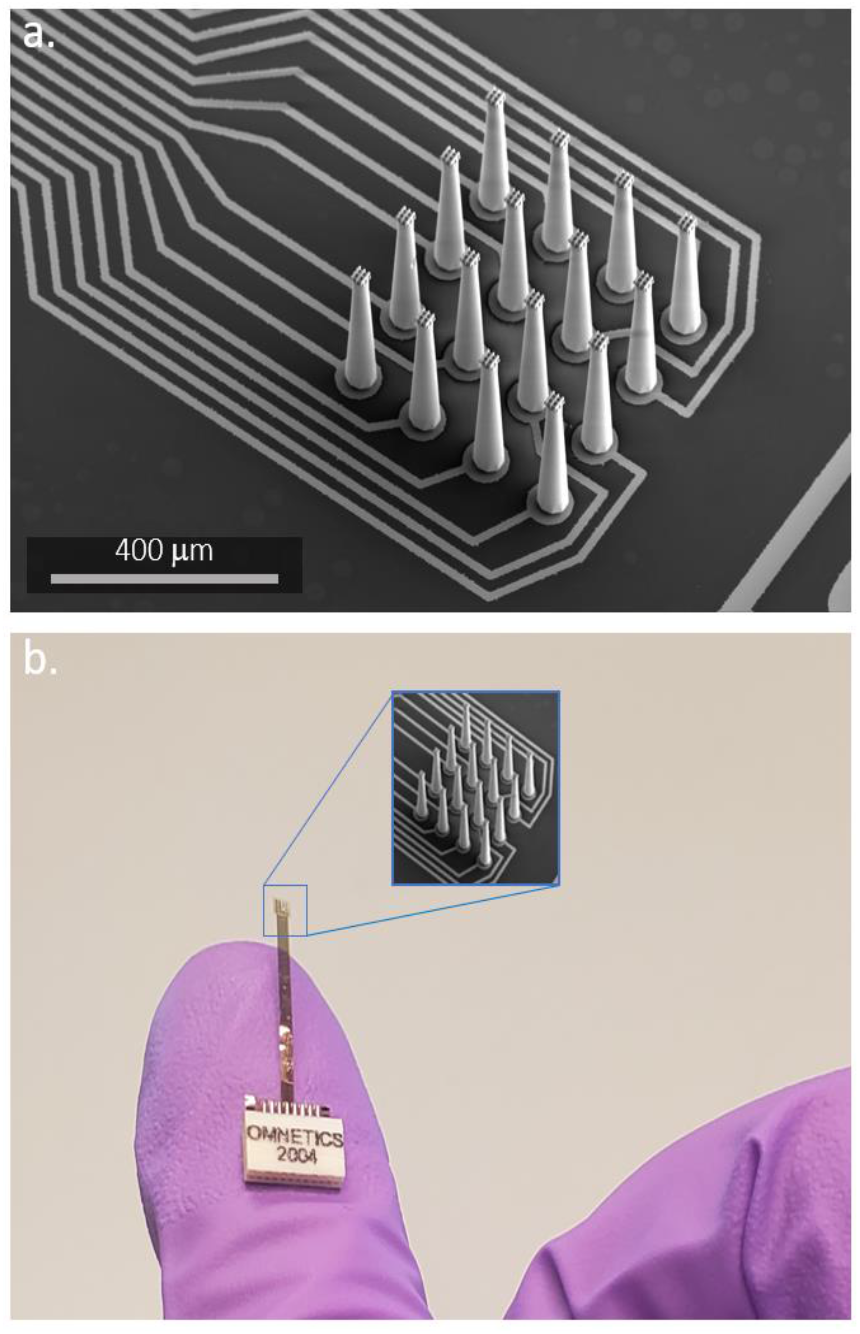
16-channel electrode array. a, SEM micrograph showing traces and 3D printed electrodes fabricated via direct laser writing. **b,** Assembled device showing polyimide flex cable, finger for scale.

**Figure 2.**
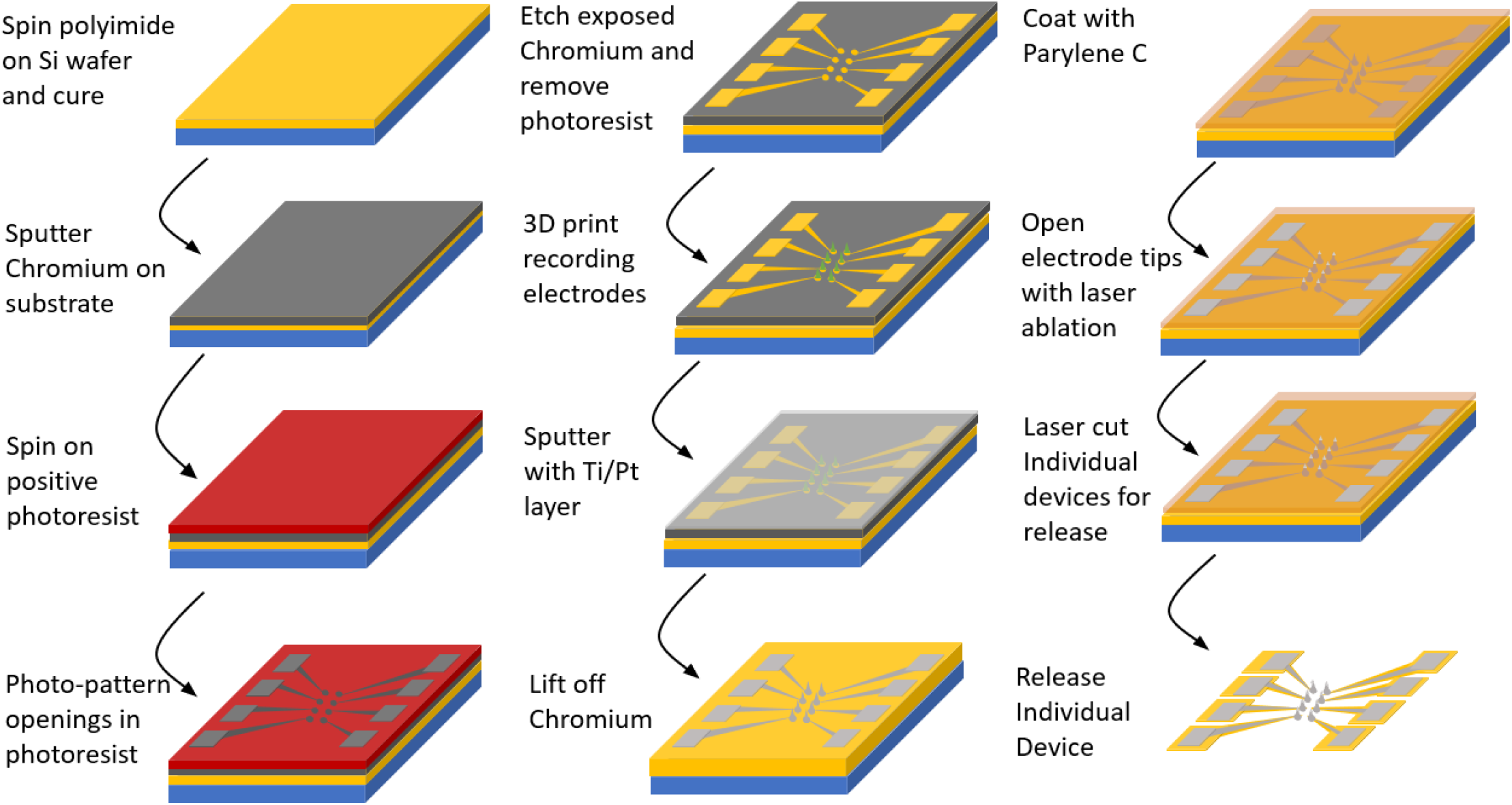
Fabrication process for recording electrodes. A metal lift off mask is patterned over the desired substrate. Electrodes are printed, metallized, and then the mask is etched off leaving the metallized prints and traces. Next, a parylene isolation layer is added over the entire device and then this insulating layer is selectively removed from the tips of the electrodes. In a final step, the full device perimeter is laser cut with a femtosecond laser (Monaco) to release individual devices from the wafer.

To insulate the metal traces and metallized spikes, the outer surface of the wafer is then coated with parylene C via vacuum deposition, a common insulation process used for electrodes [12]. To create a recording surface at the electrode tip, a small region of the parylene C isolation layer is then removed to expose the underlying platinum. In prior reports, this has been achieved using a mask and UV laser process [13], or a focused ion beam (FIB) [7]. The FIB process is slow and costly, and single-photon laser ablation with a mask provides little flexibility for opening specific locations in complex three-dimensional structures. In our fabrication process, parylene C removal is achieved via femtosecond laser milling using a high-pulse, energy-amplified fiber laser (Monaco by Coherent, max 40 μJ.) To do this, we utilize the same resonant scanning two-photon microscope that forms the basis of the 3D printer, just switching from the 80 MHz low pulse energy laser source to the Monaco. This a-thermal milling process provides the flexibility needed to remove material at micron resolution without damaging fine 3D structures. **Figure 3** illustrates the tip opening process. The exposed metal is evident in **figure 3d**, and **figure 3e** shows a scanning electron microscope (SEM) image with false color based on energy-dispersive X-ray (EDX) chemical analysis. The blue color on the exposed tip indicates a surface presence of the element platinum.

**Figure 3.**
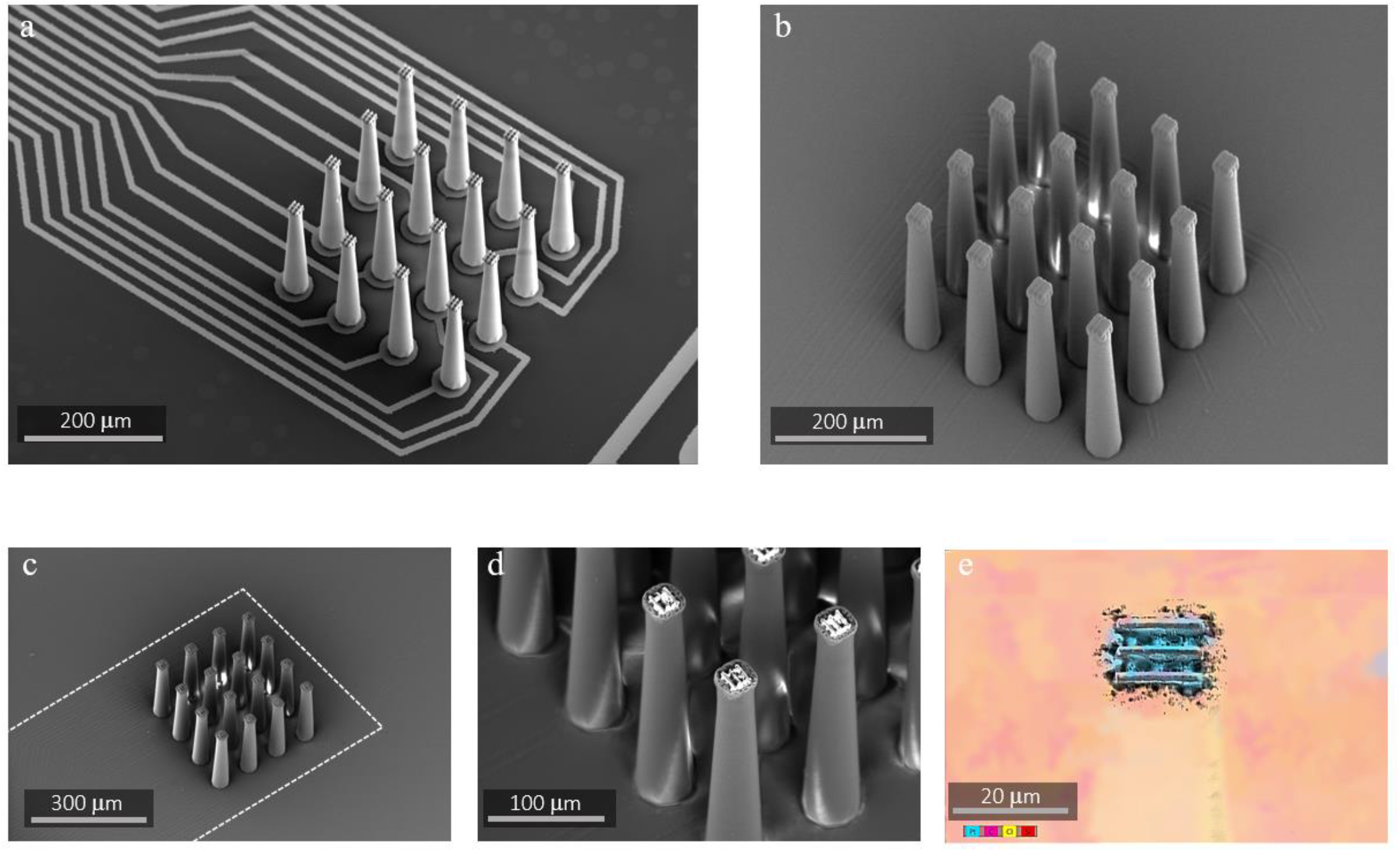
Fabrication of a 16 channel electrode for neural recording. **a,** Metallized array on wafer prior to insulation. **b,** Array coated in 3 μm of Parylene C insulation. **c-e,** Images taken post tip exposure. The connecting traces are outlined with a dashed line **(c)**. The exposed metal is evident in **(d)**, and (**e)** shows an EDX of the exposed tip with platinum shown in blue.

**Figure 4** illustrates a cross-section of the device and impedance spectra from 16 electrodes on a single test device. The mean impedance of the 16 electrodes is 200 kOhm at 1 kHz. Note that in these initial tests a “log pile” structure is 3D printed at the top of each electrode (**figure 3e**). Once metallized, this structure creates more metal below the ablation plane, increasing the metal content on the tip after ablation. (The laser ablation is not selective for parylene, and we reasoned that there could be a risk of removing too much metal without this step, but this was not examined systematically.)

**Figure 4.**
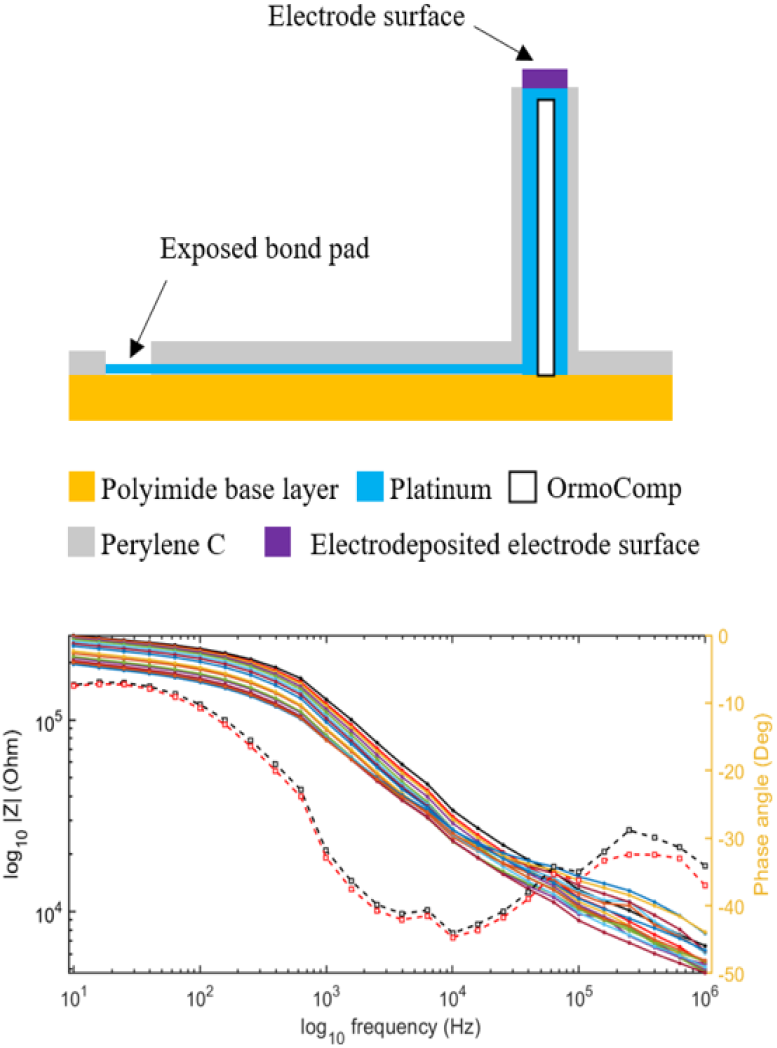
Recording surface and impedance. a, Prototype electrode cross-section. **b,** Impedance spectrum of all 16-channels from a device (filled circle)and example phase traces (open squares).

As a final fabrication step, the devices are singulated by scanning the device outline with the high pulse energy laser (Monaco) to fully cut through the polyimide. Using anisotropic conductive film (ACF), the flex cables are bonded to an Omnetics connector allowing for direct integration with Intan and OpenEphys recording systems.

The fabrication process described here provides a general recipe for high aspect-ratio 3D printed electrodes integrated on thin-film substrates. The automated printing and laser ablation steps are algorithmically defined, making it easy to alter the height profiles or shapes of electrodes and customize arrays to specific experiments or anatomical features of the brain.

### Biomimetic geometries and insertion tests

Based on Euler’s buckling calculations for the 1 GPa modulus of Ormocomp, individual electrodes with 20 μm flat tipped geometry will withstand a critical force of 1-3 mN, which is more than an order of magnitude above the penetration requirement for similar electrode shapes in another study-0.17mN [14].

However, our preliminary surgeries determined that multi-electrode arrays composed of flat topped 20 μm electrodes at a pitch of 90 microns could not be inserted in songbird brains due to dimpling of the brain surface as a result of the “bed-of-nails” effect. We found that the same 20 μm electrodes at 300 μm pitch were easily inserted in songbird brains.

To reduce tissue insertion forces [14], we looked to a geometry found in nature. The mosquito proboscis reduces insertion force while resisting buckling thanks to a tip geometry shown in **figure 5b.** With the high resolution of the 3D print process, we found that the electrode geometry can be altered to include a sharp spike resembling the point of the mosquito needle (**figure 5a**).

**Figure 5.**
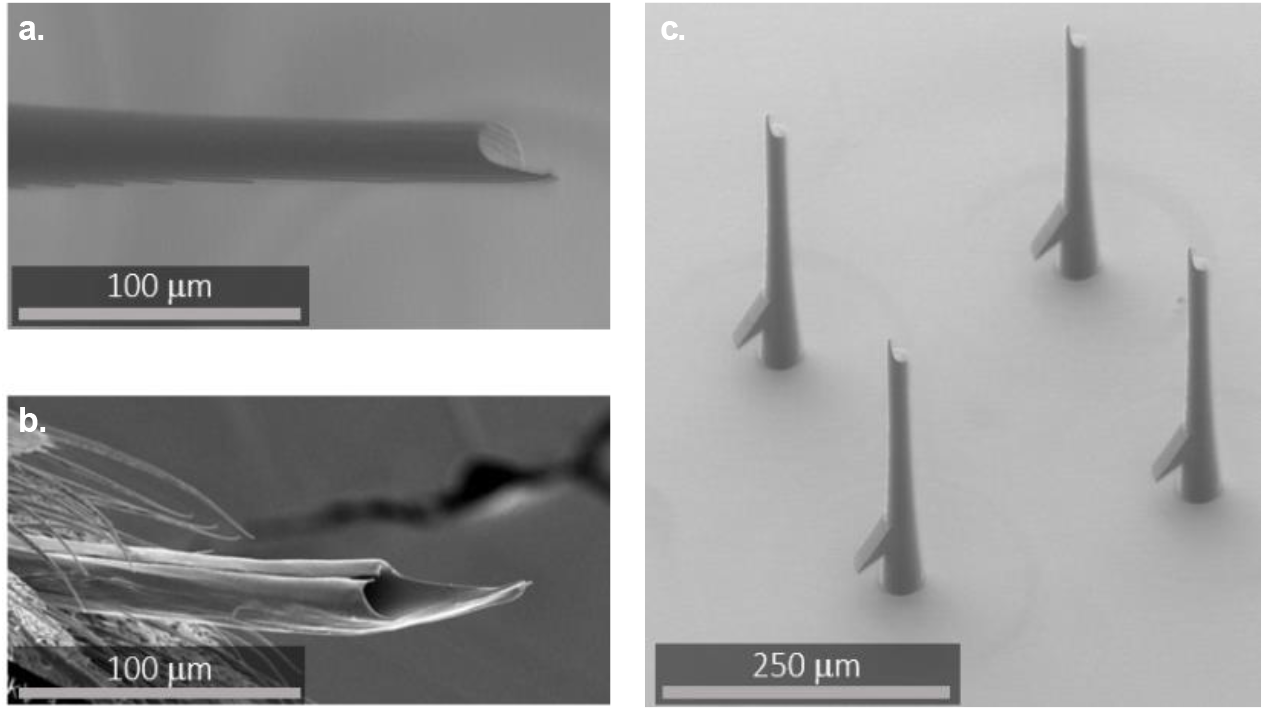
Biomimetic electrodes. **a,** Biomimetic mosquito-needle tips incorporate a sharp tip for reduced insertion force, as observed from an actual mosquito (**b,** adapted from Wikimedia Commons). **c,** These probes can be inserted through the dura of birds.

We build test structures composed of 350 um tall mosquito tip electrodes in a 4×4 grid at 150 micron pitch - a spacing where the flat topped electrodes would have caused significant dimpling of the brain. We found that the sharp tips of the biomimetic needles led to easy implant not just in brain, but even through the dura of a songbird. An implant that avoids a large durotomy will be faster and reduce the risks of swelling and damage to the brain surface. This mosquito needle design is a proof of concept only. To date, we have not built electrically connected devices with the mosquito needle geometries. This will require depositing an insulating layer over the electrodes, and then selectively removing that layer while preserving the sharp tip shape. Other sharp tip profiles such as cones or pyramids may prove to be more practical, since those shapes can be programmatically defined in the laser cutting step that removes the parylene.

### Porous stimulating electrodes

In addition to neural recording, microfabricated electrodes produced using thin-film lithography have been proposed for high-channel-count neuromodulation. Applications include bioelectric medicine via the stimulation of peripheral nerves [15, 16], visual prosthesis via retinal interfaces [17], and cortical interfaces to provide neuromodulation for mental disorders or enhance stroke recovery [18,19]. Multiple companies are now pursuing neural interfacing through microfabricated polymer electrodes, and these projects include future applications in neural stimulation [2] [20] [21]. However, as the charge-injecting surface area decreases, adequate currents for stimulation require higher voltages leading to accelerated delamination and degradation of stimulating electrodes [22–24].

**Figure 6.**
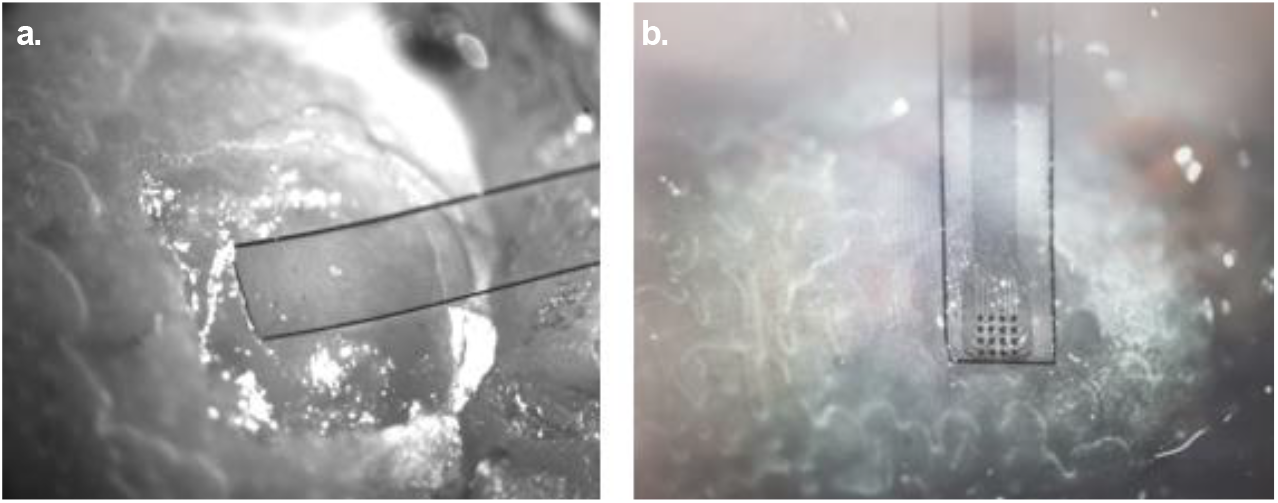
Bird implants. **a,** Insertion of test structures in a bird brain. **b,** insertion of full electrodes. Electrodes with biomimetic needle tips can be inserted through the dura at 150 μm pitch. Note the transparency of the polyimide in both images.

Traditional microfabricated electrode arrays are by nature planar. One approach to improving their stimulation capacity is to create 3D structures that are raised above the electrode surface, allowing for a more intimate connection with the target tissue. This can have the effect of concentrating charge delivery to target neurons, improving both stimulating thresholds and specificity. This concept was investigated in a silicon retinal interface with stimulating electrodes composed of vertical pillars, 10 μm in diameter and 65 μm tall, that interfaced directly with the inner nuclear layer of cells in the retina [25]. This intimate contact facilitates concentrated charge injection for retinal prosthesis and allows a lower stimulating voltage – both of which could lead to a higher resolution visual prosthesis with lower power requirements for stimulation [26].

**Figure 7.**
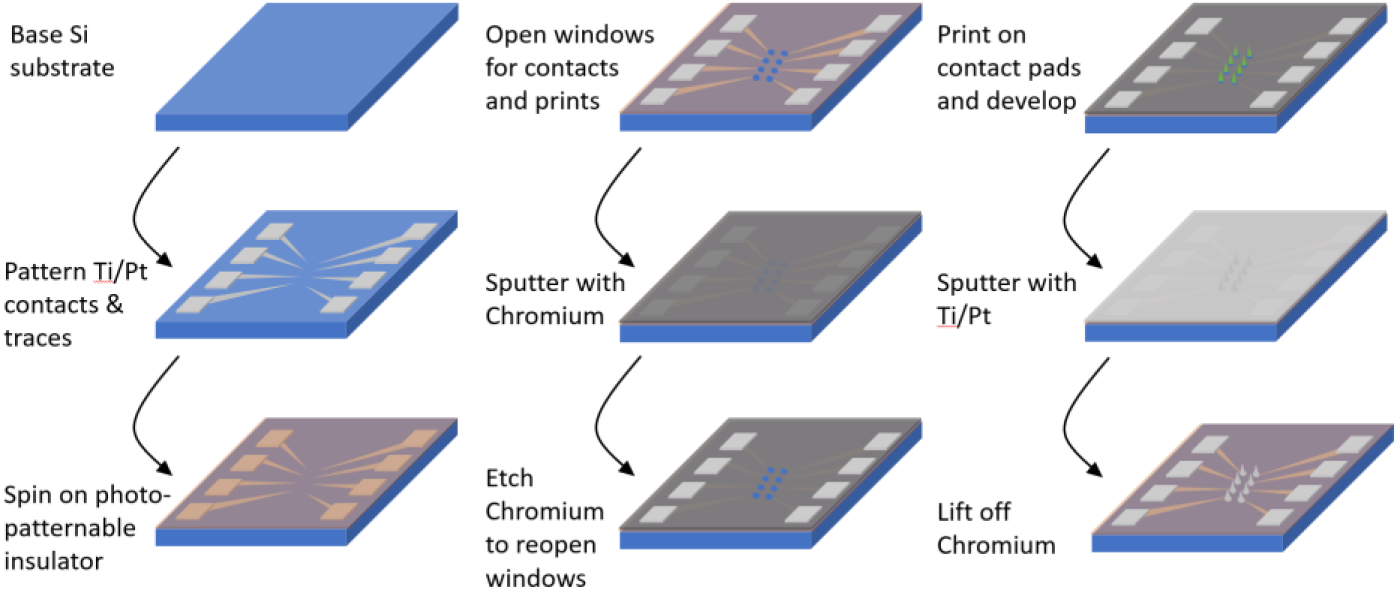
Photolithography process for Si supported devices. The metal contact pads and traces are patterned directly on the Si wafer. The traces are then isolated with SU8 and coated in a Cr etch mask, leaving an opening for prints and the contact pads through both layers. Electrodes areprinted, metallized and then the chromium is etched away leaving the metallized prints and pads.

While 65 μm tall vertical pillars on rigid silicon are an appropriate form factor for small, localized retinal implants, devices that interface with a larger fraction of the retina or other curved surfaces will require integration of stimulating pillars on a flexible substrate. Similarly, some applications such as cortical stimulation may require higher aspect ratio stimulating electrodes for therapeutic charge delivery. We anticipate that the thin-film 3D electrodes described in this project could be adapted to a stimulating electrode geometry. As a first step toward that goal, we developed a process to create macro-porous 3D electrodes. These devices were initially fabricated on silicon wafers for easy electrochemical testing.

**Figure 7** illustrates the fabrication process for the porous electrodes, based on a Si substrate. (The metal lift-off process described previously in **figure 2** involved fabricating electrical traces at the same time that the 3D prints were coated in metal. In contrast, this process involves pre-established traces that become connected to raised 3D metal surfaces in the metal sputtering step.) We printed structures with pore cross-sections ranging from 40 μm^2^ to 400 μm^2^ as well as solid prints and planar electrodes lacking 3D prints for controls. To further reduce directionality, sputtering at elevated pressure was used to accomplish interior metallization of the porous structures.

**Figure 8** shows SEM micrographs of the structures. These prototypes feature metallization throughout the interior of a porous 3D shape. **Figure 8c-d** shows the extent of the internal metallization of a print both before and after FIB sectioning. A complete device can be seen in **figure 8a**. For the shape with the largest internal pores, initial results revealed a ~2x increase in charge storage capacity relative to a flat 2D electrode pad (**figure 8b)**. Further tests will be needed to systematically compare the charge injection capacities of electrodes with different geometries and porosity parameters.

**Figure 8.**
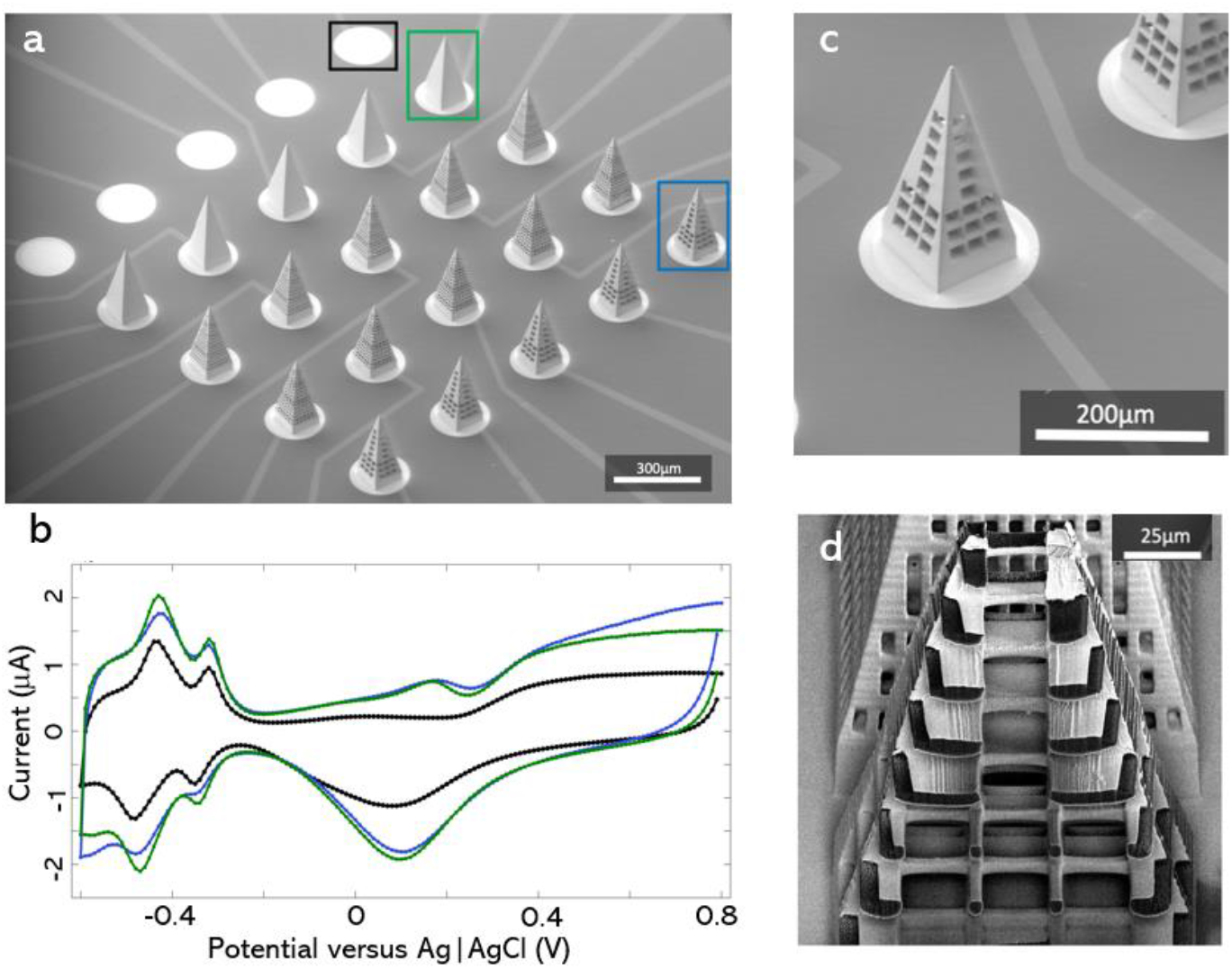
Stimulating electrodes. **a,** Metallized electrode array on a Si substrate with SU8 isolation of traces only. **b,** Example CV curves reveal increased charge injection for 3D pyramids (green and blue) relative to flat circular pads (black). Blue corresponds to a pyramid with 20 micron pores, and green a solid pyramid with no pores. **c-d,** SEM images of a print before and after FIB sectioning.

### Adhesion of 3D printed polymers to the substrate

A prevalent failure mode of these devices is the detachment of the 3D print from the polyimide substrate. Our preliminary data from sonication tests indicates that when devices are printed at a starting height precisely localized to the polyimide surface (within a 1 μm tolerance), the 3D prints do not detach from the substrate when subjected to the maximum power (ultrasonic effective power 330 W) of our benchtop sonicator. The strength of the attachment is further enhanced with the addition of the metal and insulating layers over the top. While we do not anticipate short-term print detachment or breakdown of the device, the performance of the material stack in chronic implants over longer timescales remains an important question. We have two approaches to reducing the risk that 3D printed electrodes will detach from the polyimide in chronic implants. In the first approach, adhesion was increased using laser pitting, where the photopolymer was mechanically anchored through the creation of clean, a-thermal trenches in the polyimide (**figure 9**). In the second approach, through-holes or vias were placed under each electrode spike and each electrode was finished with a backside 3D print. This effectively clamps the polyimide substrate within a covalently bonded block of OrmoComp. This method was used in our prior work developing a peripheral nerve interface [27].

**Figure 9.**
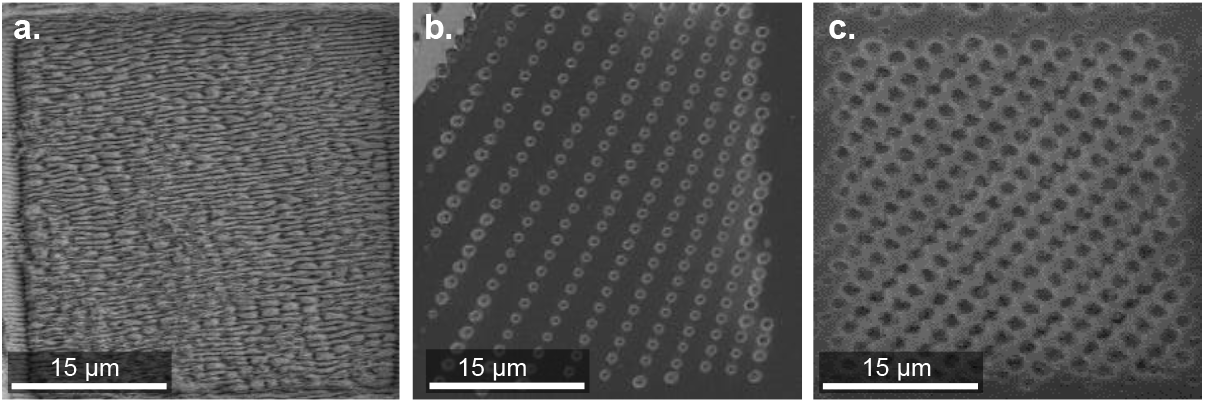
Polyimide roughened with femtosecond laser prior to printing. **a,** SEM micrograph showing roughened polyimide. **b-c,** Holes of varying density/pattern through polyimide.

### Material limitations

The prototype devices describe here include some features that will need to be changed in near term iterations of the method. For example, in the current 3D recording device (**figure 1-3**), the top and bottom layers of insulator are composed of two different materials, parylene and polyimide, without an adhesion layer between them. A titanium oxide adhesion layer will be added in future fab rounds. If delamination between parylene and polyimide proves to be a problem that can’t be adequately solved with an adhesion layer, the mixed material stack could be replaced by a single material composition of parylene top and bottom layers. Similarly, the current electrode uses a titanium layer above and below the platinum to bond the metal to the polymers. Delamination of this polymer/metal interface is a common failure mode for thin film electrodes, particularly at the top surface where polymer is being deposited on an oxidized layer of titanium [28]. In future iterations, this top and bottom adhesion layer could be replaced with silicon carbide to provide a superior adhesion layer between polymers and metal. In the longer term, we would like to replace the parylene outer insulating layer with silicon carbide (deposited with plasma enhanced chemical vapor deposition) – both to improve barrier properties and to increase stiffness of electrode spikes. This stiffer insulating layer would allow some reduction in the minimum electrode diameter that can be implanted without buckling.

## Discussion

The goal of this project was to create 3D electrodes with flexible geometry, fabricated at the highest resolution possible. This emphasis on maximizing resolution in 3D-printed electrodes is motivated by multiple factors. First, the achievement of high resolution and design flexibility in two-photon lithography opens a path for unique designs and a wide range of electrode shapes. Biomimetic mosquito-needles that penetrate through dura, barbs that allow electrodes to anchor within tissue, and pores in electrodes that could promote charge injection or tissue ingrowth are all possible with our high-resolution two-photon based fabrication methods. The 3D printing process enables not only unique electrode geometries, but also customization of electrode length profiles to match the curvature or depth profiles of specific brain regions.

A second benefit of high-resolution 3D printing is that it enables a reduction in electrode cross-section which in general will lead to chronic neural recordings with higher signal to noise ratio (SNR). For electrodes that are significantly larger in cross section than a neuron, a reactive tissue response encapsulates electrodes and damages cells at distances up to 100 μm from the implant [29–33]. Action potential amplitude decays rapidly with distance from an electrode [34–36]. As a result, adverse tissue response is a particularly severe problem for small animal studies where a large encapsulation area will prohibit single neuron resolution recordings from the densely packed regions of interest. The two-photon lithography employed here will allow fabrication of microelectrodes with dimensions well below the 20 μm limit that is thought to evade much of the brain’s immune response [37–39].

Finally, increased resolution enables the fabrication of high-channel-count electrode arrays, resulting in a greater number of electrodes per unit area within the brain, or retina. These devices represent a region of electrode configuration space that was previously unoccupied, namely the potential for high-channel-count 3D electrode arrays with features definable at micron resolution.

### Small animal model electrodes

We anticipate initial applications in small animal models where 3D electrode arrays could be fabricated to conform to specific spatial profiles within target brain regions. In addition to chronic implants in fixed locations, integrated flex cables allow these devices to be mounted on micro-drives to sample multiple depths serially. In songbirds and other small animal models, the quality of single unit isolation decays after implantation, making it challenging to record many single units in learning-focused studies that evolve over time. This progressive loss of signal is a major motivation behind our electrode design work, because the only known way to improve signals over the long run is to reduce electrode scale. While carbon fiber electrode arrays have demonstrated stable recordings over long time scales, yield is low and the fabrication process is not scalable [40]. Thin-film silicon carbide ultramicroelectrodes provide a scalable alternative that will be investigated further [41], and polymer electrodes using insertion shuttles show promise for increasing signal longevity [42]. Until the progressive loss of signal is resolved, the highest signal to noise ratio recordings will be achieved by moving multi-electrode arrays into fresh brain tissue with a micro-drive. Micro-drives that advance an electrode with a screw or small motor have been the cornerstone of electrophysiological studies in songbirds and mice for decades [43], and high-channel-count silicon probes have increased the yield of these experiments [44]. Still, for a single silicon shank with many electrodes, only the electrodes near the tip of the probe are moved into fresh tissue when a micro-drive is advanced. For the 3D electrodes described here, every electrode contact will be moving into fresh tissue when the micro-drive is advanced. We anticipate that high-channel-count 3D electrodes on micro-drives will yield a greater number of single unit recordings, extend the operational period of animal experiments, and provide the unique ability to sample 3D volumes in behaving animals.

### Porous stimulating electrodes

We anticipate multiple benefits from porous stimulating electrodes. At the simplest level, macro-pores can potentially increase the surface area of the electrodes while maintaining the same overall displaced tissue volume. For example, the solid pyramids in **figure 7a** have a surface area of 0.076 mm^2^ while the pyramids with the smallest pores have a total surface area of 0.391 mm^2^, providing a five-fold increase in the contact area between the stimulating surface and the tissue. The benefit of this porosity remains to be determined for chronic implants where tissue ingrowth leads to increased access resistance within the pores. While tissue ingrowth may be a limiting factor for charge-injection, it could potentially stabilize the porous neural interface against micromotion and reduce fibrotic encapsulation. Ingrowth of neural processes could also be encouraged to promote “neurotrophic electrodes” with the potential for stable multi-unit recordings and reduced thresholds for stimulation of ingrown axons or dendrites [45–47]. Recent studies have demonstrated vascular integration of porous ECoG arrays implanted at the surface of the brain in mice [48]. We speculate that similar vascular ingrowth could occur in porous, 3D-printed microelectrodes and that this could improve device longevity by providing stabilization against migration and micro-motion.

As a separate benefit, the protruding surfaces of 3D stimulating electrodes will provide better electrical contact between the electrode and neural tissue, with potential applications in cortical micro-ECoG recording and stimulation, as well as peripheral nerve interfacing [26].

Achieving chronic electrical stimulation requires microelectrodes with a sufficiently high charge injection capacity. For intracortical stimulation using conventional microelectrodes with geometric surface areas of 500-2000 μm^2^, the charge thresholds for eliciting an electro-physiological response is approximately 1 nC/phase and tissue damage can occur at approximately 4 nC/phase [49,50]. Our prototype electrodes are platinum-based and inject charge via surface-confined faradaic reactions and double-layer capacitance. In future tests we anticipate further enhancements of charge injection capacity via sputtering or electrodepositing high charge injection micro-porous coatings such as Iridium Oxide onto the surface and interior of the 3D prints.

## Conclusion

The fabrication method for 3D-printed electrode arrays presented here is robust, wafer scale, and fully compatible with standard Si and flexible polyimide device fabrication processes. Using high-resolution 3D laser writing, a wide range of unique electrode shapes ranging from biomimetic needles to porous electrodes can be fabricated. These devices will enable volumetric recordings at a spatial resolution not achievable with current 3D microelectrode devices. We anticipate the method providing new tools for a range of users across neuroscience and neural engineering research, and ultimately in human applications such as visual prosthesis or nerve interfaces where high density recording and stimulation are required in a small form factor.

## Methods

### 3D printed electrodes

Three-dimensional electrode structures are printed using a two-photon 3D printer, in a process known as direct laser writing. This printer was described previously [11]. Electrode shapes are designed in standard 3D CAD software and uploaded into the custom PrintImage software as .STL files for print voxelization. The photopolymer used is a hybrid resist based on the commercially available photoresist OrmoComp^®^ (Micro Resist Technology), a glass-like, biocompatible member of the Ormocer family [51]. We add a photo initiator, ((2,4,6-trimethylbenzoyl)phosphine oxide (TPO), Sigma Aldrich), a stabilizing agent, (3,5-Di-tert-butyl-4-hydroxytoluene (BHT), Sigma Aldrich) and fluorescein (Sigma Aldrich) for *in situ* imaging during 3D printing. A 780 nm Chameleon Discovery laser with a 100 fs pulse width, 80 MHz rep rate, and ~40 mW power through a 20x Nikon immersion lens (NA 0.7) is used to initiate polymerization in the photoresist. Following prints, the substrates are submerged in Ormodev (Micro Resist Technology) developer for 12 hours to remove un-polymerized photoresist, followed by a rinse in isopropanol. The development process is followed by a 10 min. UV cure at 395 nm (Solis-365 C at 2.8 mW/mm^2^) to increase the overall degree of crosslinking in the polymerized resist and to enhance the mechanical stability of the structures [52].

An overview of the basic fabrication process can be seen in **figure 2**. Electrodes are fabricated on prime grade 75 mm Si wafers with 300 nm of thermal oxide (University Wafer). A base layer of polyimide (HD Micro Systems PI2611) is spun onto the surface and cured at 350°C for 30 min. in a nitrogen environment to a final thickness of 6 μm. Adhesion promoter (HD Microsystems VM652) is added to the edge of the wafer prior to the polyimide. 500 nm of Cr is then sputtered onto the polyimide surface (3 mTorr DC) as a sacrificial mask layer. To define the metal traces of the electrode array, AZ-1512 photoresist (Kayaku Advanced Materials, Inc.) is spun onto the wafer surface and patterned using a Süss Microtec MJB4 mask aligner (350 W mercury arc lamp) with an exposure dose of 70 mJ/cm^2^. All photolithography masks are written in-house on a direct write laser lithography system. After development (AZ 300 MIF), room temperature Transene Chromium Etchant 1020 (ceric ammonium nitrate/nitric acid) is used to etch through the 500 nm Cr layer, forming the mask for the final traces and defining the print locations. The photoresist is then removed via sonication in acetone at 37kHz for 5 min.

Prior to DLW 3D printing, the patterned wafer is then cleaned with oxygen plasma for 90 sec in a March plasma etcher at a pressure of 300 mTorr and RF power of 100 W to increase print adhesion. Following initial wafer alignment, surface finding and printing for each electrode in the array proceeds in an automated fashion. Following print and development of 3D electrode structures, the devices are plasma cleaned again prior to final metallization to increase metal adhesion to the prints. The final print structures are then sputtered with Ti (15 nm) / Pt (200 nm) in an Angstrom Engineering sputter system at 3 mTorr (Ti – DC) and 10 mTorr (Pt – RF). Lift-off of the Cr sacrificial layer is done in Transene Chromium Etchant 1020 at 60°C for 15 min. To effectively remove all metal flakes, wafers are transferred to multiple fresh etchant baths during the lift-off process with agitation, followed by a thorough rinse in DI water. Finally, the Omnetics connector contact pads are masked with Kapton tape and a 3 μm thick layer of Parylene C is deposited (Labcoater 4200) over the wafer.

The porous electrodes illustrated in **figure 8** were fabricated directly on the Si wafers as described in **figure 7**. Here an initial metallization (10/50 nm Ti/Pt) layer was patterned and then electrically isolated with 500 nm of SU-8 negative photoresist (Kayaku Advanced Materials, Inc.). Openings in the SU-8 were made using a photomask and an exposure dose of 65 mJ/cm^2^. These openings were large enough to include a region of the Ti/Pt metal traces to create electrical connections in the subsequent metal sputtering. 500 nm of Cr is then sputtered onto the polyimide surface (3 mTorr DC) as a sacrificial mask layer. AZ-1512 photoresist (Kayaku Advanced Materials, Inc.) is spun onto the wafer surface and patterned to define the print location holes in the chromium aligned with the holes in the SU8. Patterning of this layer is done with a Süss Microtec MJB4 mask aligner (350 W mercury arc lamp) with an exposure dose of 70 mJ/cm2. After development (AZ 300 MIF), room temperature Transene Chromium Etchant 1020 (ceric ammonium nitrate/nitric acid) is used to etch through the 500 nm Cr layer, forming the openings for the final print locations. The photoresist is then removed via sonication in acetone at 37kHz for 5 min. Devices were then printed, metallized, and the chromium layer released as described above.

#### FIB milling

Focused ion beam (FIB) milling is utilized to slice open printed structures and assess the extent of internal metallization. A Thermo Fisher Helios FIB was used with an argon/O_2_ beam.

#### Device finalization and release

A 1035 nm wavelength pulsed laser (Coherent, Monaco) is used to remove the Parylene C from the tips of the prints. This laser is co-aligned with the 3D printing laser (Discovery) in the system described above. The initial alignment points on the wafer are found using the Discovery to avoid damage. The Parylene is then removed from the tips of all 16 electrodes in a single cut process at a 1 MHz pulsing setting. In order to release the full device from the wafer, the 1035 nm pulsed laser is used to cut through the polyimide and Parylene layers using programmed motion of a precision translation stage. The wafer is then placed in warm water to release the individual devices. Finally, Omnetics connectors are attached to the device pads via anisotropic conductive film (3M, ACF 7371).

#### Electrochemical measurements

Cyclic voltammetry (CV) and electrochemical impedance spectroscopy (EIS) data was collected using a high surface area Pt counter electrode and an Ag/AgCl reference electrode on a Gamry Reference 600 potentiostat. Measurements were conducted in a phosphate buffered saline (pH ~7.2) consisting of 0.126M NaCl, 0.081M Na2HPO4 and 0.022M of NaH2PO4. Prior to measurement, the electrolyte solution was sparged with He gas for ~30 minutes to remove dissolved O2. CV curves were cycled at 100 mV/s until differences between subsequent scans were no longer observed. Additional full device checks were performed via Open Ephys using Intan chips.

## Acknowledgements

We thank the staff at the University of Oregon Technical Sciences Administration for assistance with hardware design as well as the facilities and staff at the Center for Advanced Materials Characterization in Oregon. This work was supported by the Phil and Penny Knight Campus for Accelerating Scientific Impact and by the University of Oregon Initiate of Neuroscience, the NIH fellowship grant F32MH118724 (M.A.B), and NIH grants R01NS104925 and R01NS118424 (T. Gardner, co-PD/PI).

